# The effect of a canine semen activator supplementation or addiction on the long-term refrigeration quality of dog spermatozoa

**DOI:** 10.1101/2021.04.09.439141

**Authors:** Marcelo Martínez-Barbitta, Claudio Rivera Salinas

## Abstract

Within modern biotechnology, different tools have been developed to maximize canine semen conservation protocol to optimizing reproductive results and making their handling more flexible. In the last decades, the survival of refrigerated semen has been prolonged from 2-3d with the first basic diluents, to 10-14d using the most modern extenders. However, their main limitation is that sperm quality decreases during cold storage. Semen activators (SA) have been produced to provide the molecules necessary to maximize the sperm survival and quality with the aim to enhance fertility and prolificacy. In this study, the effect of SA was recorded by daily evaluation of chilled semen 14d. For this experiment, six adult healthy Neapolitan Mastiff dogs, were used as donors and the semen was manually collected. Spermatozoa-rich fractions of each suject was chilled using a new generation extender for long periods of time (d0) starting from the d1 to d14, different aliquot, with (experimental trial) and without SA (control trial), were evaluated daily for motility vigor, morphology and membrane integrity. The initial sperm concentration of extended semen was 417.3±170.4×10^6^/mL (mean ± SEM) with 85.89±4.76% of MNS (morphologically normal sperm), 84.47±5.22 % vital sperm and a pH of 6.2±2.8. The initial vigor was 3.83±0.48, but after one min with SA, it rose to 4.45 ± 0.45 (P<0.001). The semen motility parameter increase significantly (P<0.05) in experimental trial, respect to control, starting to d2 at finish (except for d7). The vigor analysis significantly increase in experimental trial (P<0.05) during the most day of the study with the exclusion of d3 and d14. For evaluate the semen characteristics over time, the experiment was divided into T1 (d0-d5), T2 (d6-d10) and T3 (d11-d14) (P<0.001) in evaluation of morphology and membrane stability. The MNS reached 70% at d10 and finally 65% at d14, being considered normal and possibly fertile. With Host-s, 65% of MNS were also achieved at d14. The presence of glucose and fructose in the diluents used for refrigeration can exert very important effects given the fact that metabolic routes have been found in both sugars, providing both different and complementing effects. It can be concluded that the use of SA prior to artificial insemination improves the quality of chilled semen significantly, although it does not reverse the effects of deterioration due to cellular metabolism over time.

## 1. INTRODUCTION

Despite of the artificial insemination in dogs is documented, for the first time, by Lazzaro Spallanzani in 1788, only in the last decades its use in companion animals reproduction has been more widely performed. In fact canine pure dog breeders increasingly require the use of this tecnique using fresh, chilled or frozzen semen. The use of fresh sperm is indicated in male or female that cannot the natural mating for physical or behavioural causes. The storage of sperm (chilled or frozzen) allows a wider genetic improvement of breed advantage of chilled respect to fresh semen are cost relatively inexpensive between countries, animal transportation, and less stressing life-threatening and time-consuming reduces venereal disease risks and allows breeders to use semen from genetically superior dogs, simplify and popularize insemination techniques [1-2].

During the storage process of semen, for limited the damages caused by drop temperature, and to provide energy, maintain pH and osmolarity, reduce oxidation, preserve plasma, acrosomal and mitocondrial membrane integrity, etc., an appropriate diluent must be used (extender) with sperm [3-7].

Storage of refrigerated semen at 4-8 °C induces a transition in the sperm plasma membrane from the cooled crystalline to the gel phase. At body temperature, the metabolism of the sperm is maximum, while at room temperature (24-29 °C), it decreases. For every 10 °C decrease, cellular metabolism is reduced by 50%; at 5 °C metabolic activity of sperm is only 10% of what it would be at body temperature [8].

More recently, use of supplementation of activators of insemination (SA) whose basic composition is formulated by easily metabolized carbohydrates that provide the mitochondria of the spermatic neck with a fast energy substrate to maximize their metabolism at the time of insemination has been studied by several authors [9-12] and enhances seminal motility in order to maximize fertility and prolificacy of semen.

Therefore, use of long-term refrigeration seminal diluents together with activators when using the germplasm, would enhance the management of these biotechnologies by the reproductive male, allowing the female to be managed more effectively, maximizing reproductive results [11-14]. Therefore, the objective of this experiment was to evaluate the motility and survival of the spermatozoa under a dilution protocol with refrigeration for 14 days, corroborating the effect of SA during the whole process.

## 2- MATERIALS AND METHODS

### 2.1 Animals and location

The work was carried out in the month of December, at the facilities of the laboratory and semen bank of Clone^®^ Chile (Santiago de Chile).

The experiment was carried out according to the bioethics and animal experimentation standards of the participating countries and has been evaluated by the corresponding committees. Six Napolitan Mastiff male dog with an average weight and age of 88 ± 12 kg (MED ± SEM) and 30.5 ± 3.5 months respectively were used. The dogs were confirmed healthy based on history, clinical examination including full andrological evaluation and ultrasound examination of the prostate and testis. Dogs were fed twice daily (at 8.00 AM and 10.00 PM) using commercial dry food and selected chicken prey. Fresh water was available ad libitum.

### 2.2 Semen collection and evaluation

Semen was manually collected as described previously [15] in the same day arround 8:00 PM and promptly examined the characteristic of seminal fluids, under a laminar flow chamber (Biobase^®^ BBS-H1500, Chile). In particular the volume of the sperm rich fraction was determined, the pH of the seminal and prostatic fraction was assessed by pHmetro (Hanna Instrument^®^ HI 5521, Chile), sperm morphology and vitality were assessed by eosin-nigrosine staining (Minitüb^®^, Tiefenbach, Germany) smears counting at least 200 spermatozoa per slide, and total sperm concentration was obtained by photometer (SDM 1 Photometer, Minitüb^®^, Tiefenbach, Germany, Series 1260162485, calibrated for canines) in accordance with the manufacture recommendations.

Moreover in the sperm rich fraction, the evaluation was performed, with two aliquots of 30 µL on slides, covered with coverslips, all pre-tempered at 37 °C with thermal stage (HTM-MiniTherm, Hamilton^®^ Thorne Biosciences -Beverly, MA, USA), for visualization using a phase contrast microscope (Olympus^®^ BH-2 -Japan) X 100, waiting 60 s for observation, to determine motility (%) and vigor according to the scale described by Howard et al. (1990) [16]. The evaluation was repeated at five, 10 and 15 min.

Within the first minute of collection to a second aliquot of semen 30 µL was added 30 µL of SA extender (TherioSolutions^®^ Canine AI extender – Chile -SA-), homogenized and waited 60 s to reassess motility and vigor, as described above. The evaluation was also repeated at five, 10 and 15 min.

From the first day and daily since, the structural and vital characteristics of the semen samples were evaluated after tempering for 60 s by the vital morphology stains of Farelly (Minitüb^®^, Tiefenbach, Germany), using the method described by Oettlé (1986) [17] X 1000 with immersion oil.

Sperm vitality due to membrane stability, counting at least 200 sperm cell per sample, was performed using the Hyposmolarity Test (Simplified Host –Host-s-), according to the method of Sánchez & Garrido (2013) [18].

In both fresh and refrigerated semen, sperm with a functional membrane were considered, which reacted to hypo-osmotic stress by dilating the distal part of the spermatic tail or curling it. While those sperm without changes in the tail were considered functionally damaged, and the results were expressed as a percentage of sperm with a functional membrane (sperm with abnormalities in the tail, by Farelly, were excluded from the count).

The individual semen samples were refrigerated at 4 °C (Bozzo^®^ Refrigerator SD-350, Chile, with a maximum and minimum thermometer inside) and evaluated in relation to their vigor and motility every 24 hours for 14 days (TherioSolutions^®^ Canine Chilling extender, Chile). The alcohol in the external tube prevented cold shock during the chilling process and temperature variations over study period [19].

### 2.3 Statistical Analysis

The mean and SEM of spermatozoa vigor, motility, structural morphology and permeability membrane were measured by using descriptive statistics. The data were analyzed by ANOVA and T-test using SPSS^®^ 21.0 software (SPSS Inc., Chicago, IL, USA) [20]. The difference between values was considered significant when the P value was less than 0.05.

## 3-RESULT

### 3.1 General Characteristics

The collection of the spermatozoa of the animals showed a good seminal quality determined for a race of giant individuals during the summer season. The volume of the second fraction of the ejaculate was 3.9 ± 1.6 mL with a pH of 6.2 ± 2.8. The prostatic fraction maintained the same value and dispersion in pH. The sperm concentration was 417.3 ± 170.4 million sperm per mL. The initial evaluation of the semen by eosin-nigrosin staining resulted in 85.89 ± 4.76 with no head staining, considering them to be alive.

### 3.2 Analysis of Seminal Vigor

The initial vigor was 3.83 ± 0.48. After 1 min of AS, the vigor was 4.45 ± 0.45 (P <0.001). Graph N°1 shows the variation in the time of the refrigerated seminal vigor after being adjusted at 37 °C with and without the addition of AS.

**Graph N° 1 -.**
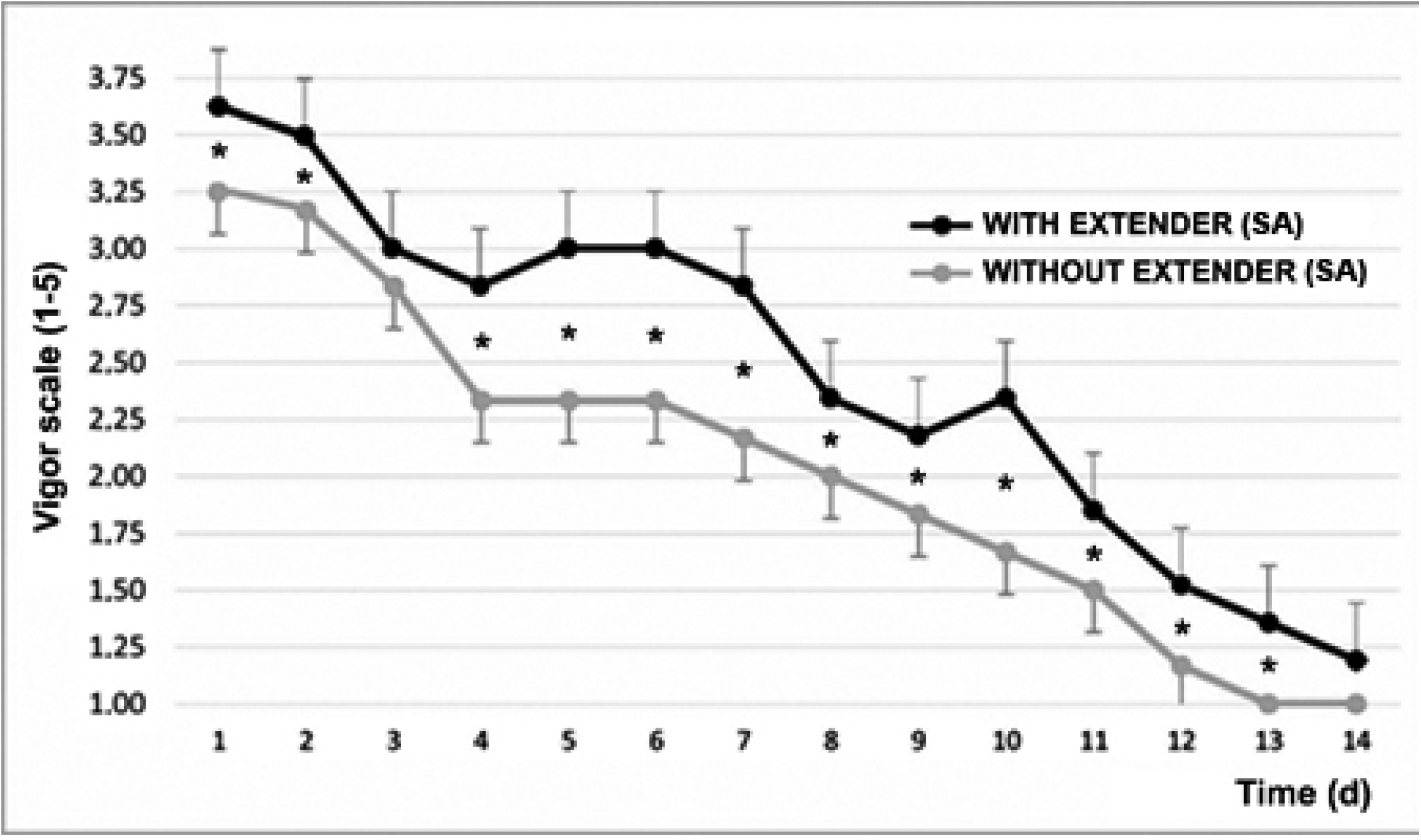
Variation of seminal vigor (according to Howard, et al. 1990) [16] after cooling and timing1. Mean ± SEM * P <0.05 ^1^The graphed evaluations correspond to 1 minute after adding the semen activator.

From d1 to d13 they showed significant differences, with the exception of d3 (P <0.065).

### 3.3 Seminal Motility

The initial seminal motility was 85.83 ± 5.23%. After one minute of AS, motility increased to 89.92 ± 5.18% (P <0.05). Graph N°2 shows the variability of seminal motility with and without AS during the days of refrigeration of the semen. At all moments evaluated, with the exception of d1 and d7, significant differences were found between the samples.

**Graph. N° 2 -.**
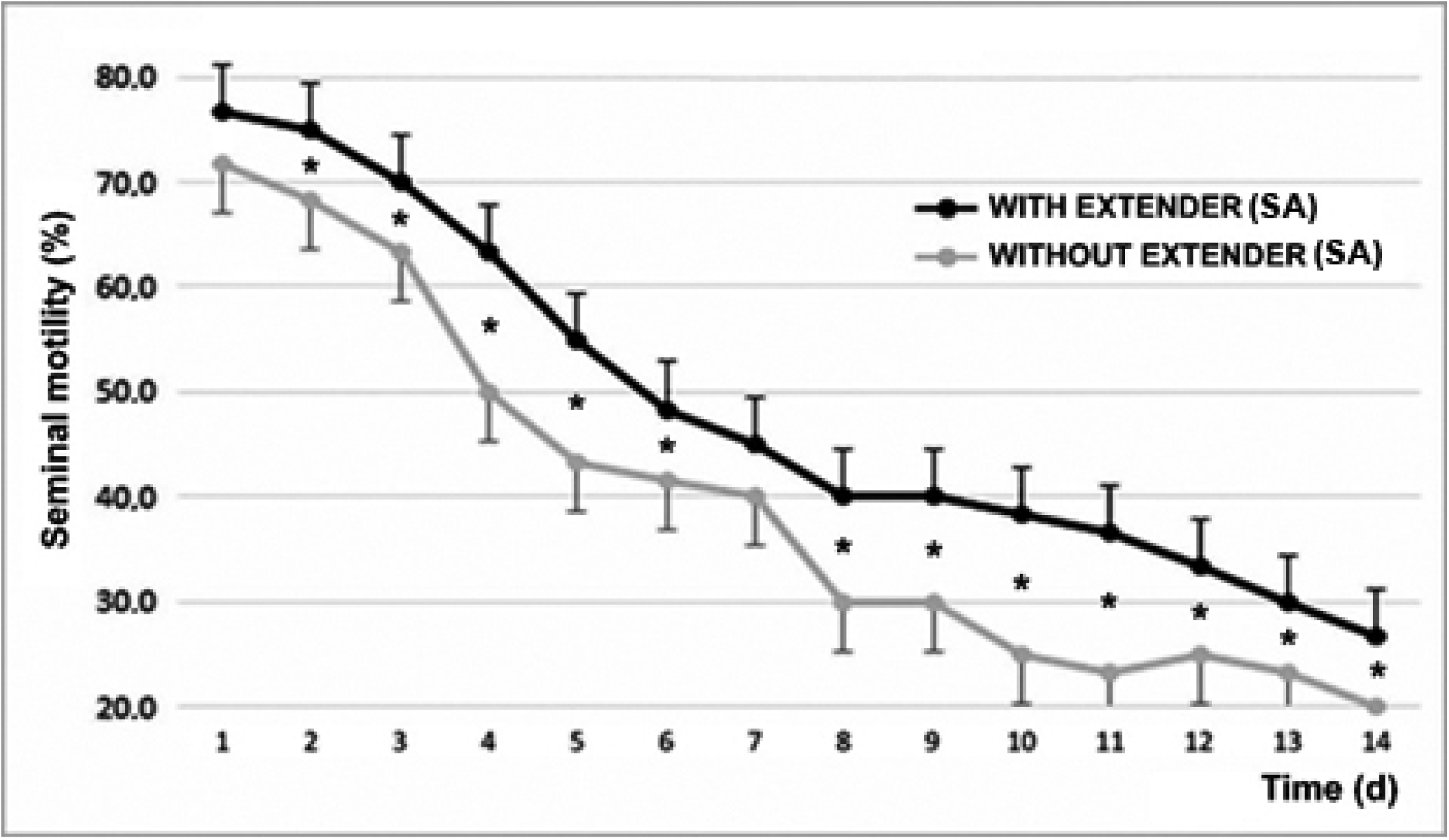
Motility with and without the effect of seminal activator during the 14 d of the evaluation of refrigerated semen^1^. Mean ± SEM * P <0.05 ^1^The graphed evaluations correspond to 1 minute after adding the semen activator.

In Graph N° 3 the variability of the average seminal vigor and motility of the entire period is observed, based on the differences according to the waiting of the evaluation, after the temperament of the sample refrigerated at 37 °C.

**Graph. N° 3 -.**
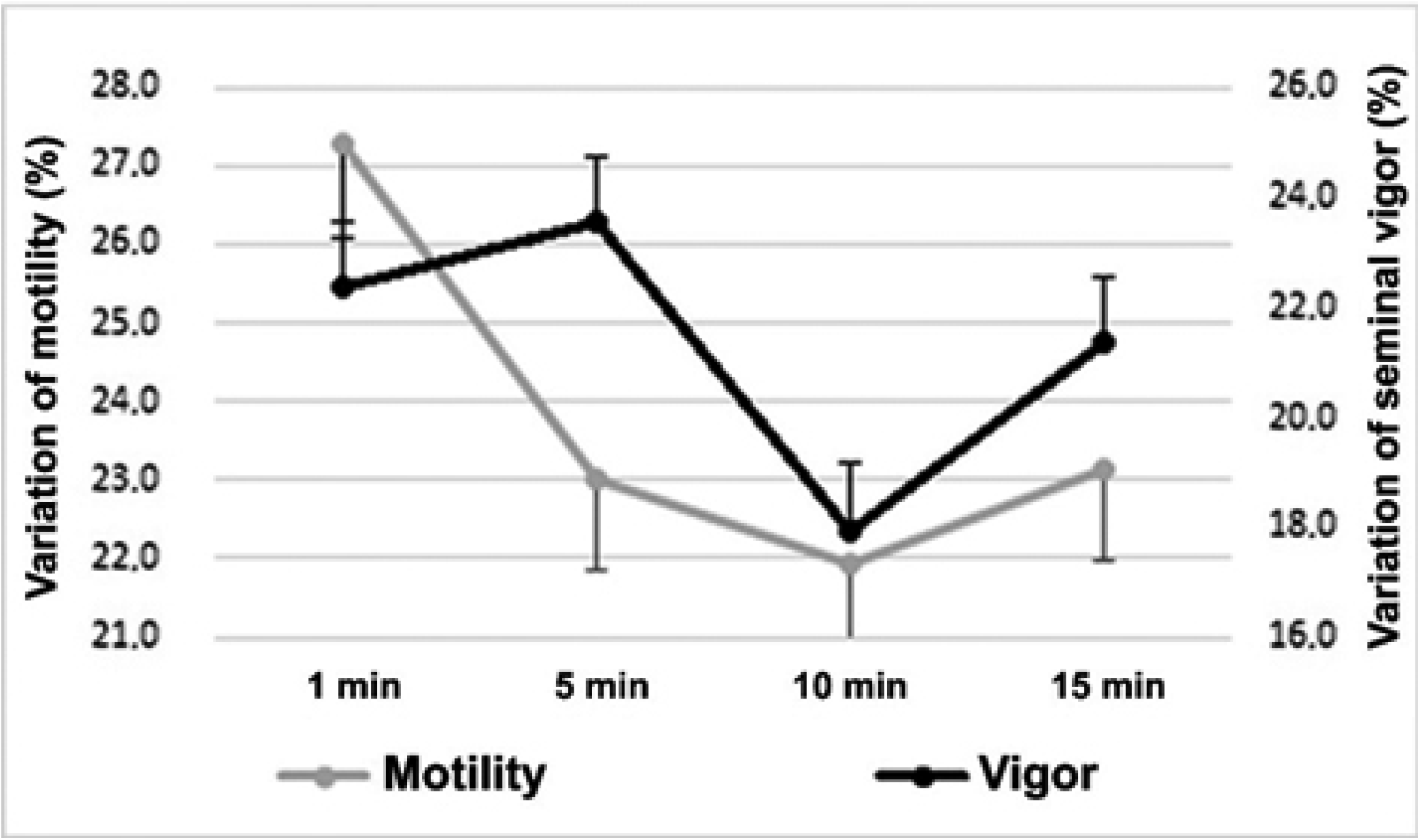
Percent variation of vigor and seminal motility average of 14 days, after the placement of the semen activator (Mean ± SEM)* *The variation was calculated as the difference between each value (motility and vigor) of each day with the course of the evaluation time (1, 5, 10 and 15 minutes).

### 3.4 Spermatic Morphology

Regarding morphology, 84.47 ± 5.22% of MNS (morphologically normal sperm), 3.96 ± 1.95 with head defects, and 7.20 ± 3.01 of defective intermediate piece were found with vital staining and finally 4.37 ± 0.61% of tail defects at the beginning of the experiment. In Graph N° 4 that observes the evolution of the defects and their distribution in the spermatozoa during refrigeration.

After 14 days of decrease, with a practically linear decrease, a 19.47% lower value was found, with an average reduction of 1.5%/day, according to the Farelly staining analysis method. It should be noted that distinguishing between the affected area, tail changes increase by 49%, while in the intermediate part they increase by 122% and finally, problems in the sperm head grow by 216% (P <0.001).

**Graph. N° 4 -.**
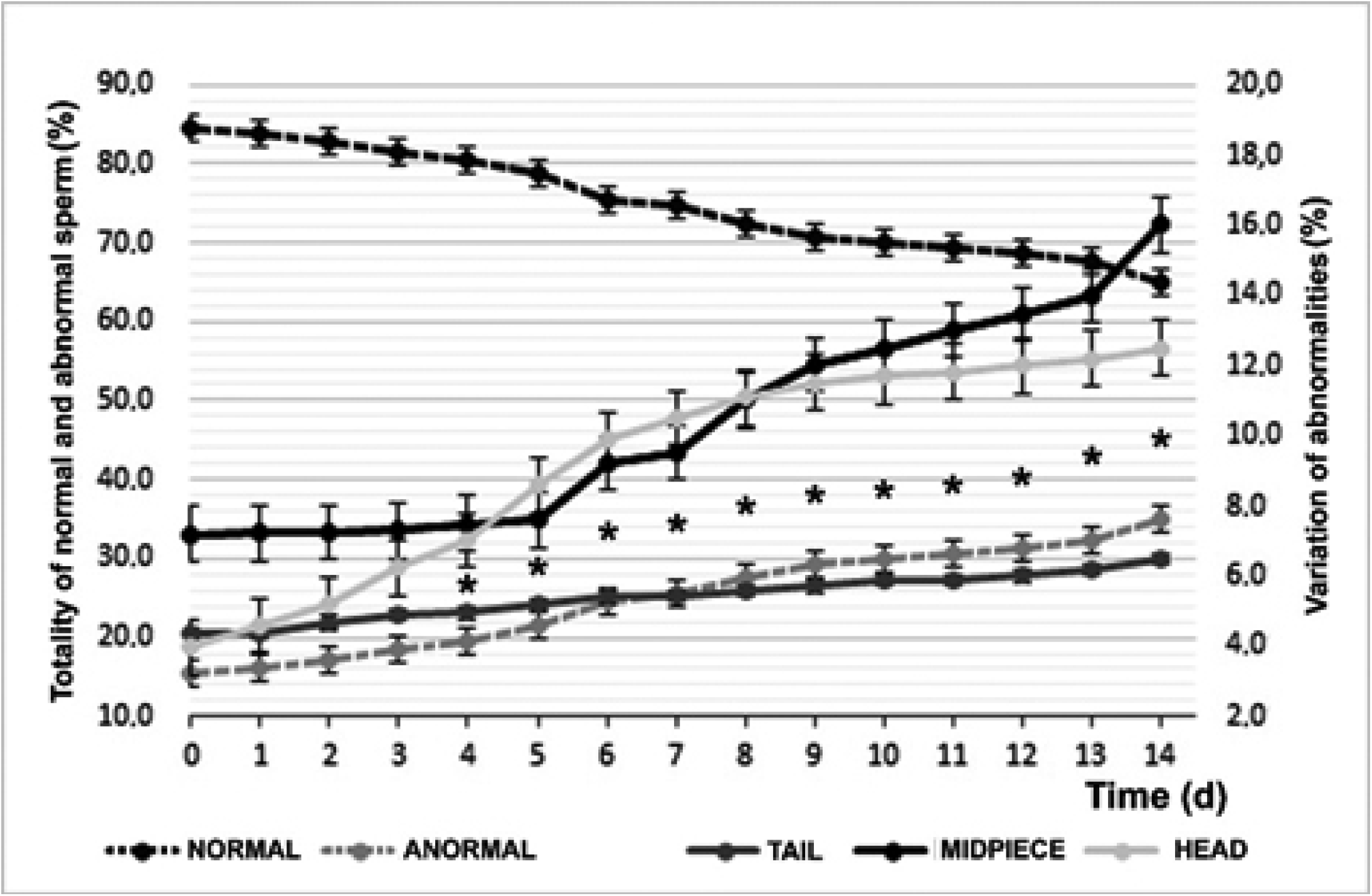
Temporal variation of sperm morphology by vital staining (Mean ± SEM). *Difference between abnormalities of head or middle piece and significant tail (P <0.05).

According to the waiting period of the evaluation of the motility and vigor of the semen samples, after the placement of the SA, there were very important changes. At one minute of placing the SA, the average vigor during the 14 days was 2.43 ± 0.65, and at 15 minutes it increased to 3.23 ± 0.86 (P <0.01), being 32.8 % higher. For motility, the variation was less but still substantially equal (P <0.05), while the average evaluation of the period one minute after adding the SA was 48.45 ± 12.9%, at 15 minutes it was 56, 55 ± 15.11%, an increase of 16.7% at the end of the term.

### 3.5 Membrane Integrity

Regarding the stability of the membrane, at d0 85.89 ± 4.76% of spermatozoa were found to be normal, subtracting 14.11 ± 4.76% of those with permeability integrity defects. In the Graph N° 5 shows the variability of sperm membrane stability.

**Graph. N° 5 -.**
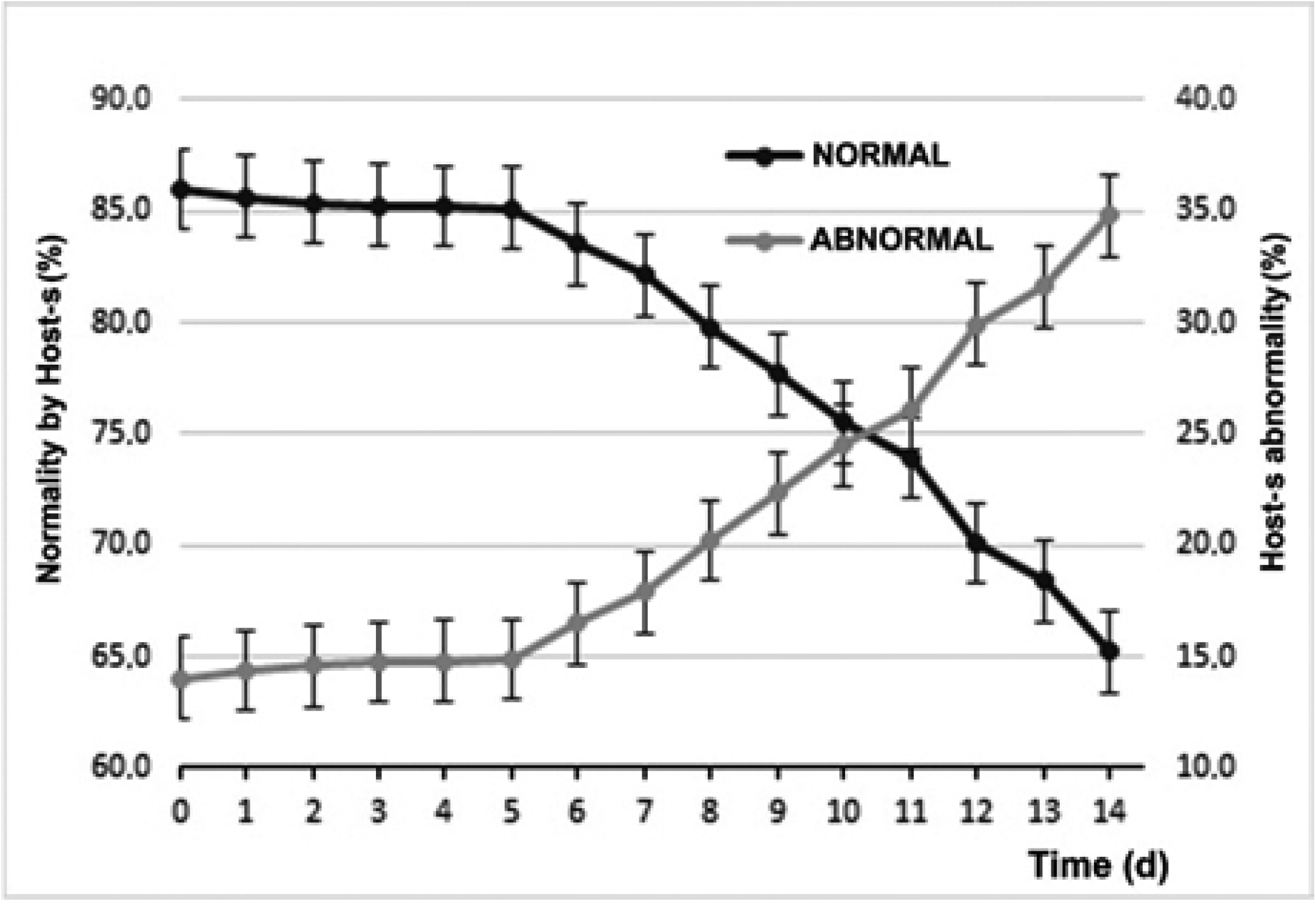
Temporal results of refrigerated semen Host-s samples^1. 1^Those termed ABNORMAL were those that did not undergo morphological changes after Host-s.

To analyze the changes in membrane stability and sperm morphology, the times were divided into three periods, according to the viability of the seminal extensors that are normally divided into short, medium and long-term. Thus, T1 (short term) is used for the period between d0 and d5, T2 (medium term) for the period between d6 and d10 and finally T3 (long term) between d11 and d14. In the worksheet N° 1 are the variations relative to the changes in seminal morphology and membrane stability.

**Table N° 1 -.** Comparison between sperm morphology and membrane stability test with distinction of storage times by refrigeration. T1 = d0-d5; T2 = d6-d10; T3 = d11-d14.

|  | T1 (%) | CV (%) | T2 (%) | CV (%) | T3 (%) | CV (%) |
| --- | --- | --- | --- | --- | --- | --- |
| **Normal Morphology<sup>1</sup>** | $81,97 \pm 0,89^a$ | 2,67 | $72,62 \pm 1,06^b$ | 3,28 | $67,62 \pm 0,94^c$ | 2,78 |
| **Membrane Stability<sup>2</sup>** | $85,43 \pm 0,13^d$ | 0,38 | $79,71 \pm 1,45^e$ | 4,06 | $69,40 \pm 1,81^f$ | 5,22 |

<sup>1</sup> Refers to Farelly's Vital Staining subjective assessment.
<sup>2</sup> Corresponds to normal sperm according to Host-s.
Different letters in the same row $p < 0.001$

## 4 – DISCUSSION

The volume, concentration, live and dead sperm (eosin/nigrosin) and pH of the second phase of the ejaculate of all animals remained within normal values, therefore they were all included in the following analysis [21].

Regardless of species, all extenders used for cold storage of genetic material must provide cells with: nutrients as an energy source; a buffer against damage by changes in pH; electrolyte concentration to control physiological osmotic pressure; protection against bacterial growth and thermal shock during refrigeration and/or freezing [3-4,7]. By contrast, activators are added agents, used in both refrigerated and frozen semen (although also with fresh semen), to enhance sperm activity and mainly have the function of providing rapidly available sugars for sperm cells as well as providing a buffer against intense metabolic activity that would imply an associated decrease of seminal longevity and intrinsic fertility [22]. They can also integrate other substances with preventive or conservative activities of the product itself or beneficial ones associated with semen handling (for example, antibiotics). The objective of semen storage using chilling procedure is to preserve gametes at low temperatures, without reaching the freezing point, which induces deleterious intracellular changes that affect viability and fertilizing potency of spermatozoa [15,22]. This technique of chilling preservation is a promising alternative to conventional semen cryopreservation, and is easily adapted for clinical use. It is particularly useful for the shipment of semen where the cost of procedures and materials are high [23]. This technique would also be of major importance if it extended the fertilizing potential of spermatozoa for more than a few days (ideally between 10-14 d). Thus semen could be collected a few days after the detection of pro-estrus in bitch and then processed, transported and stored at the location of its intended use.

The commercial formulations of the laboratories reveal known components, but also have components that are not disclosed in the formulas made by the manufacturers. TherioSolution^®^ declares that it possesses glucose extender in SA extender and chilling products and also fructose in the chilling formula. Most of the extensors reported for use in cryopreservation of domestic canine sperm contain either glucose, lactose or fructose [15,19,23-32]. Furthermore, fructose has been used intensively in extenders for wild dogs such as foxes [24-27,33]. Sugars have long been included in semen diluents as exogenous energy substrates, as osmotic components, and as cryoprotective agents [34]. Sperm can glycolyze glucose, fructose and mannose [35] and oxidize arabinose [36]. However, the higher molecular weight of sugars such as lactose, sucrose and raffinose have low permeability, and are generally considered only as a cryoprotective agent [37] and not as an energy source. In dog semen, due to the absence of seminal vesicles, fructose is found in a very low concentration compared to that of other species [30,38]. However, canine sperm appear to be able to use fructose in a similar way to sperm from species with high fructose content in their accessory gland secretion. Still, it is important to notice that in 2001 Iguer-ouada & Verstegen found better results using glucose versus fructose as the energy substrate, possibly indicating a preference of dog sperm to metabolize this molecule instead of fructose [23].

Ponglowhapan, et al (2004) [30] investigated the importance of the source and concentration of the type of energy source molecule in sperm metabolism during refrigeration in canines and found that lactose is more efficient than glucose in obtaining energy levels higher (ATP measurement enzymatically) in fresh sperm. Furthermore, there are indications that fructose may possibly play a role as a sperm activator after ejaculation [39]. Yildiz, et al. (2000) [40] suggest that sugars allow the maintenance of osmotic pressure and perform a cryoprotective action. Based on this, is expected that the inclusion of sugars in the extenders could serve as the preservation of the spermatic motility, maintenance of the viability, and integrity of the acrosome and the spermatic membrane [40-41].

During our evaluation, in 14 d, either the partial results of the use of chilling extender using motility and vigor as indicators of vitality and Host-s as a test of plasma membrane stability, showed very promising results. And at almost all times, the addition of the SA extender improved the quality of the evaluated sperm, proving that this SA extender could be added at any time to stimulate quality and improve reproductive results. In the 14 days of evaluation, the MNS barely reached 70 % at d10, reaching 65% at d14. These were able to be considered normal values that would not affect the fertility achieved with said semen [21].

A high variability was found between the motility and vigor values achieved according to the evaluation time after the placement of the SA (Graph N°3). From one to 15 minutes, the values always increased and had significant variations, which could be explained by the absorption of the seminal plasma membrane of the activator extender components over time, which takes some time and increases the metabolic activity of the sperm cell. It also establishes the importance of waiting time between the addition of the activator and the evaluation of the sample. This proves the necessity to be strict with the time frame of the evaluation and the methodology of the applied experimental design.

The use of Host-s, for hypo-osmotic stress or membrane stability, has been validated in dogs for both fresh and refrigerated semen [18], in which the capacity of the membrane of the sperm cells to allow the flow of ions and of water into the cell is measured [42]. Our results showed a stability up to d5 of 85%, then it decreased almost linearly until d14 reaching 65%, decreasing approximately 2% per day. It was very interesting that after two weeks, two out of three sperm evaluated, remained with a viable plasma membrane. Previous studies [43] had already shown that the addition of sugars significantly differentiated the response of sperm against hypo-osmotic tests in control samples in the absence of sugars. Reported that the motility of sperm cells had a higher significant value with the use of refrigeration media with monosaccharides (glucose and fructose), compared to disaccharides (trehalose and sucrose). This could be explained by their greater availability, either as an energetic medium for the spermatic neck mitochondria as well as their cryopreservative function.

The results could be explained because during semen refrigeration, the main function of sugars is to provide the energy substrate required by sperm for the normal performance of their functions, glucose being one of the sugars best used by the sperm cell [41]. Fernández-Novell et al. (2004) [44] observed in dog sperm that glucose, but not fructose, can specifically activate the protein kinase AKT, involved in the regulation of several important cellular metabolic processes. This would imply that glucose would directly activate all AKT-regulated pathways of sperm. It can be assumed that, in the case of the canine, sugars can act not only as proper substrates of metabolism, but also as direct modulators of spermatic function. The effects of these two sugars on the metabolism of freshly ejaculated sperm have been studied in dogs, and there is evidence that dog sperm metabolizes glucose and fructose using separate pathways [41]. This results in differentiated management systems of energy as indicated by their different roles in glycogen metabolism [45], motility patterns [39], hexose metabolism [41], and glycogen deposition [46].

Fructose, with respect to glucose, showed an increase in speed in the metabolic pathways and, therefore, in the formation of ATP. Fructose, by increasing the motility-related consumption rate of ATP, results in a faster and more linear specific pattern than that observed with glucose [39], dedicating most of the energy consumption of the sperm to maintaining motility. The effect of fructose on motility, would be related to a strong increase in the phosphorylation index of hexoses with respect to glucose. This increase, in addition to the consumption of ATP in the tyrosine phosphorylation, could lead to the establishment of the substrate that completes the cycle in which the energy that is immediately lost is generated. A drop in intracellular ATP levels would logically induce an immediate increase in ADP, which in turn would activate the glycolytic rhythm and increase ATP formation. When ATP returns to high levels, there would be a simultaneous drop in ADP levels, with a consequent decrease in glycolytic rhythm [47].

This feedback phenomenon would cause a high consumption rate and possible depletion of the fructose provided in the diluent, which indicates the importance of choosing the appropriate type and concentration of sugar, since small variations could cause large changes in the functional state, and therefore, the capacity of survival of sperm stored in refrigeration. Hence, the proposal analyzed by technicians and researchers to renew refrigeration diluents to maintain the metabolic activity of sperm and delay their death. In turn, the presence of both monosaccharides (glucose and fructose) is also highlighted in the extender for chilling used, where each monosaccharide would act differently on seminal cell metabolism, but only glucose in the SA extender.

Mammalian sperm require exogenous substrates for a variety of functions. For example, to preserve intracellular energy stores, cellular components, and most importantly, to support motility [48]. They can obtain energy through mitochondrial oxidative phosphorylation and glycolysis, by consuming glycolizable sugars, such as glucose, fructose, mannose, and maltose [49]. Fructose is believed to be an important source of energy for ejaculated sperm [50], and along with glucose it is found in seminal plasma in many mammalian species.

The results of the present study clearly demonstrated that the main effect of glucose and fructose on cold semen extenders in canine semen are to provide inputs that intervene in sperm motility and movement patterns. Motility is an important indicator of the use of sugar by sperm since they provide the essential external energy source to maintain motility. This is the practical criteria for evaluating semen quality on a commercial level, being widely associated with fertility [15].

There are not many studies that have published variations in seminal quality after prolonged periods [23,30]. Also, none that have used SA in each period to assess the reaction of sperm to the addition of products that enhance their activity, finding more than important and significant reactions throughout the evaluation period, which could improve the viability of the cells and their mobility, which would ultimately affect the fertility and prolificacy obtained [49].

During d1, although there were apparently no modifications in the plasma membrane evaluated by Host-s, the morphological abnormalities increased significantly between 5-6%. Within these, especially those associated with problems in the sperm head (the abnormalities found were doubled from 4 to 8%, being highly significant), motility decreased by 25% (highly significant) and vigor decreased by one point. These findings are consistent with that published by Ponglowhapan et al. (2004) [30], where these researchers found that in those first five days, the consumption of carbohydrates by the sperm cells was greater than a posteriori, which would indicate a deceleration of the metabolic rate of the sperm that could compromise the subsequent preservation and cell survival. Although possible changes in the capacitation and reactions of the sperm acrosome were not evaluated in the present work, several publications [26,31,51-53] reveal that the significance of these harmful effects in the seminal cell, which transcend cell death and lower fertility, are frequent in the freezing/thawing process, but not as pronounced in keeping the semen above freezing point.

In this work, we found that the highest values of motility and vigor were obtained between 10 and 15 minutes after the activator was placed in the semen, when related to the evaluation at the moment of placing the SA, which would demonstrate the rapid availability of carbohydrates for cellular metabolism and the use of the latter by seminal cells.

No matter how much diluent is used to protect the sperm, heat shock will cause significant cell death. In our study, without the inclusion of the activator, there was a decrease of 32.91% in motility and 41.45% in vigor, initially, between d0 and d1, without the application of SA, data in the sense of previous research [23,30].

## 5- CONCLUSIONS

Refrigeration of semen for long periods, obtaining quality semen, allows the sending of samples over long distances, avoiding the mobilization of breeders with the associated stress, transportation costs and potential health risks of mobility. The simplicity of the technique allows its mass use at a commercial level. The use of SA extender enhancers significantly improves the quality parameters at any time, after tempering the refrigerated sample, which would improve the fertility and prolificacy results of the obtained litters. These latter estimates require further studies to be verified.

The results obtained open the expectation to new working modalities with extenders for chilling that allow the use of breeding animals of high genetic value to be generalized, without the need to freeze the semen (with the dramatic changes that freezing/thawing cause in the seminal cell), with the complexities of transportation, and costs and handling that they require to obtain satisfactory results.

This research did not receive any specific grant from funding agencies in the public, commercial, or not-for-profit sectors.

## 6- CREDIT AUTHOR STATEMENT

Martínez Barbitta: Project administration, Conceptualization, Investigation, Software, Data curation, Validation, Writing-Reviewing and Editing. Rivera Salinas: Financial support, Methodology and Writing-Original draft preparation.

## 7- ACKNOWLEDGMENT

The authors thank the team of Dr.VM Héctor Pérez from SVSL Puerto Rico for their collaboration in the translation of the article.

